# Double-μPeriscope: a tool for multi-layer optical recordings, optogenetic stimulations or both

**DOI:** 10.1101/2021.06.28.450155

**Authors:** Mototaka Suzuki, Jaan Aru, Matthew E. Larkum

## Abstract

Intelligent behavior and cognitive functions in mammals depend on cortical microcircuits made up of a variety of excitatory and inhibitory cells that form a forest-like complex across six layers. Mechanistic understanding of cortical microcircuits requires both manipulation and monitoring of multiple layers and interactions between them. However, existing techniques are limited as to simultaneous monitoring and stimulation at different depths without damaging a large volume of cortical tissue. Here, we present a relatively simple and versatile method for delivering light to any two cortical layers simultaneously. The method uses a tiny optical probe consisting of two micro-prisms mounted on a single shaft. We demonstrate the versatility of the probe in three sets of experiments: first, two distinct cortical layers were optogenetically and independently manipulated; second, one layer was stimulated while the activity of another layer was monitored; third, the activity of thalamic axons distributed in two distinct cortical layers were simultaneously monitored in awake mice. Its simple-design, versatility, small-size and low-cost allow the probe to be applied widely to address important biological questions.

## Main text

Despite over half a century of intense investigation, we still do not understand the role of layers in the complicated microcircuitry of the cortex^1^. Over the last decade optogenetic manipulation – either activation or inhibition – of specific cells^2,3^ has been established as a powerful method to examine the causal relationship between the specific cell-type and the network activity, cognitive functions or animal behavior^4^. Multichannel electrical recording is the most widely used approach to measure the neural responses to optogenetic stimulation. This approach is preferred over the optical approaches available (e.g., two-photon imaging) because of the relative ease of use and lower costs. The problem with this approach, however, is that extracellular signals represent the summation of the activity of all structures (dendrites, axons, and cell bodies) from all cell-types surrounding each electrode; even worse, extracellular activity is usually very complex *in vivo*, particularly in the awake brain^5^. Polysynaptic activations can also occur through local and long-range connections.

One approach to overcome this problem is to use two-photon microscopy with two light sources – one for optogenetic stimulation, the other for imaging – and visualize how stimulation of specific cell-type evokes responses in surrounding neurons^6^. However, most approaches are either limited in range (i.e., the total depth) or speed. Moreover, it requires expensive equipment that is unaffordable for many laboratories. A more recently developed method is to optogenetically stimulate or image ~50 neurons that are arbitrarily distributed in three dimensional space^7,8^; however, the required equipment is unaffordable for many laboratories and the combination of this 3D stimulation with imaging of arbitrary chosen cells in 3D space would be even more difficult and thus far such a success has not been reported.

More invasive approaches using a large (1 mm) prism covering all cortical layers would allow simultaneous imaging to neurons across layers^9,10^; however, the significant damage caused by 1 mm prism insertion into the cortical tissue must be considered. Substantial bleeding during surgery is unavoidable and a significant number of cells are killed or damaged by the procedure^10^, suggesting the possibility that repairing and rewiring processes take place, even if the animal survives the injury. Moreover, this approach typically requires chronic surgery involving a substantial delay between the surgery and recordings. Less invasive approaches use a single tapered fiber that has multiple windows^11^ or μLED probes^12–14^; however, they are only capable of optogenetic stimulation and do not permit calcium fluorescence imaging.

A relatively standard approach is the insertion of an endoscope that with the aid of a right-angled microprism can deliver light to and receive light from a single cortical layer simultaneously^15,16^. To take advantage of the ease and flexibility of the single endoscope approach while allowing the probing of multiple layers, we developed a double-μPeriscope that is capable of optogenetic stimulation, fluorescence imaging or both of two separate cortical layers (Fig. 1a,b). Each μPeriscope consists of a 0.10 mm X 0.10 mm or 0.18 mm x 0.18 mm micro right-angled prism, a custom-designed GRIN lens, and a multi-mode optical fiber. To examine the focal light delivery from each μPeriscope, we stimulated two compartments of cortical layer 5 (L5) pyramidal neurons by inserting the double-μPeriscope with the lower one in cortical L5 and the upper one in layer 1 (L1) (Fig. 1c). We expressed light-sensitive ion channels, Channelrhodopsin 2 (ChR2)^2,3^ and yellow fluorescent protein (YFP) in L5 pyramidal neurons (Fig. 1d). A double-μPeriscope allowed us to address a wide range of neurobiological problems, three of which are described below.

**Figure 1.**
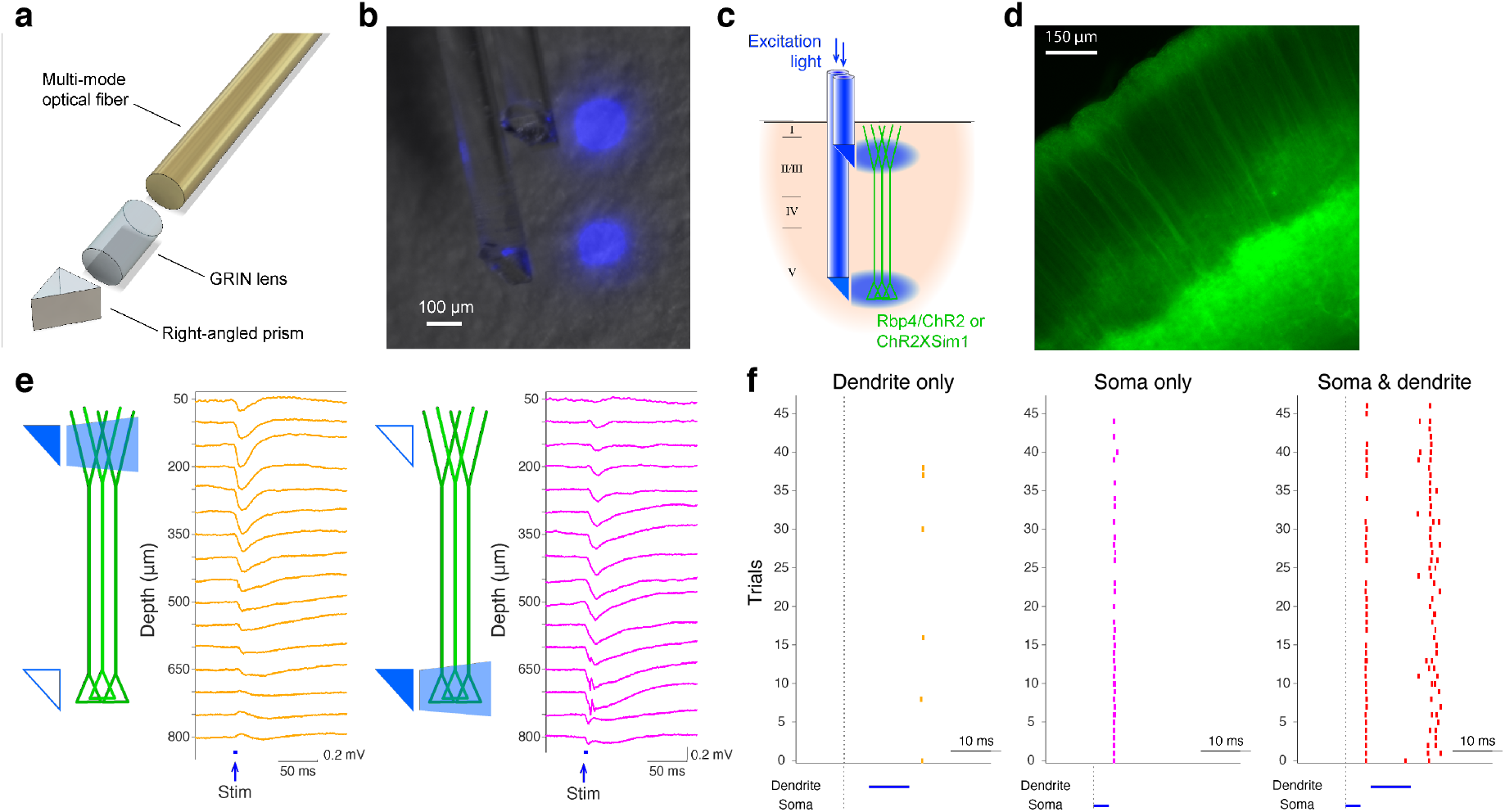
The design and functionality of double-μPeriscope. (a) An exploded view of individual μPeriscope. (b) A photomicrograph of the double-μPeriscope with two blue light spots. (c) Schematic diagram of the experiment. (d) A photomicrograph of the cortical slice where L5 pyramidal neurons express ChR2 and YFP. (e) Evoked potentials by dendritic stimulation (left) vs somatic stimulation (right). (f) Somatic APs evoked by dendritic stimulation only (left), somatic stimulation only (middle) and both (right). Vertical ticks denote APs.

First, a previous in vitro study showed that combined stimulation of cell body and distal apical dendrites within a close time window induces a large, regenerative, long-lasting calcium plateau potential that evokes a burst somatic action potentials (APs)^17^ – a phenomenon referred to as ‘backpropagation-activated calcium spike firing’ or BAC firing. However, due to technical difficulties, it was unknown whether and under what conditions BAC firing occurs in vivo. The double-μPeriscope, in combination with electrophysiological recordings from L5 allowed us to directly test this hypothesis *in vivo* for the first time. To express ChR2 in L5 pyramidal neurons, a transgenic mouse line Sim1-KJ18-Cre × Ai32 was used. In agreement with the previous in vitro data, somatic stimulation preceding dendritic stimulation *in vivo* under anesthesia evoked multiple APs, compared to the stimulation of either somatic or distal dendritic compartment alone (Fig. 1e-f), suggesting the occurrence of the backpropagating AP-induced Ca^2+^ spike firing in vivo.

The same double-μPeriscope is capable of studying the interaction between cell classes in different layers. As an example, we examined the effect of optogenetic stimulation of cortical layer 2/3 (L2/3) on L5 by expressing ChR2 and YFP in L2/3 pyramidal cells in the primary motor cortex (Fig 2a,b) through combined use of a transgenic mouse line Rasgrf2-2A-dCre with Cre-dependent adeno-associated virus vector with CaMKIIα promotor. The upper μPeriscope stimulated ChR2-expressing L2/3 pyramidal cells while the lower μPeriscope measured the calcium fluorescence in L5 where a cell-permeable synthetic calcium indicator (Cal-590 AM) was pre-injected. In agreement with the recent study in primary somatosensory area using a different approach (electrophysiological recordings)^18^, the net effect of L2/3 stimulation onto L5 turned out to be inhibitory in the motor cortex (Fig 2c,d), suggesting that the L2/3-to-L5 inhibition might be a general principle across the cortex in contrast to the prevailing textbook assumption that L2/3-to-L5 projections are excitatory^19^. The previous study suggested that somatostatin-positive interneurons are likely to be responsible for the translaminar inhibition in the somatosensory cortex^18^; whether the same mechanism mediates the translaminar inhibition in the primary motor cortex needs more investigations that are beyond the scope of this technical report.

**Figure 2.**
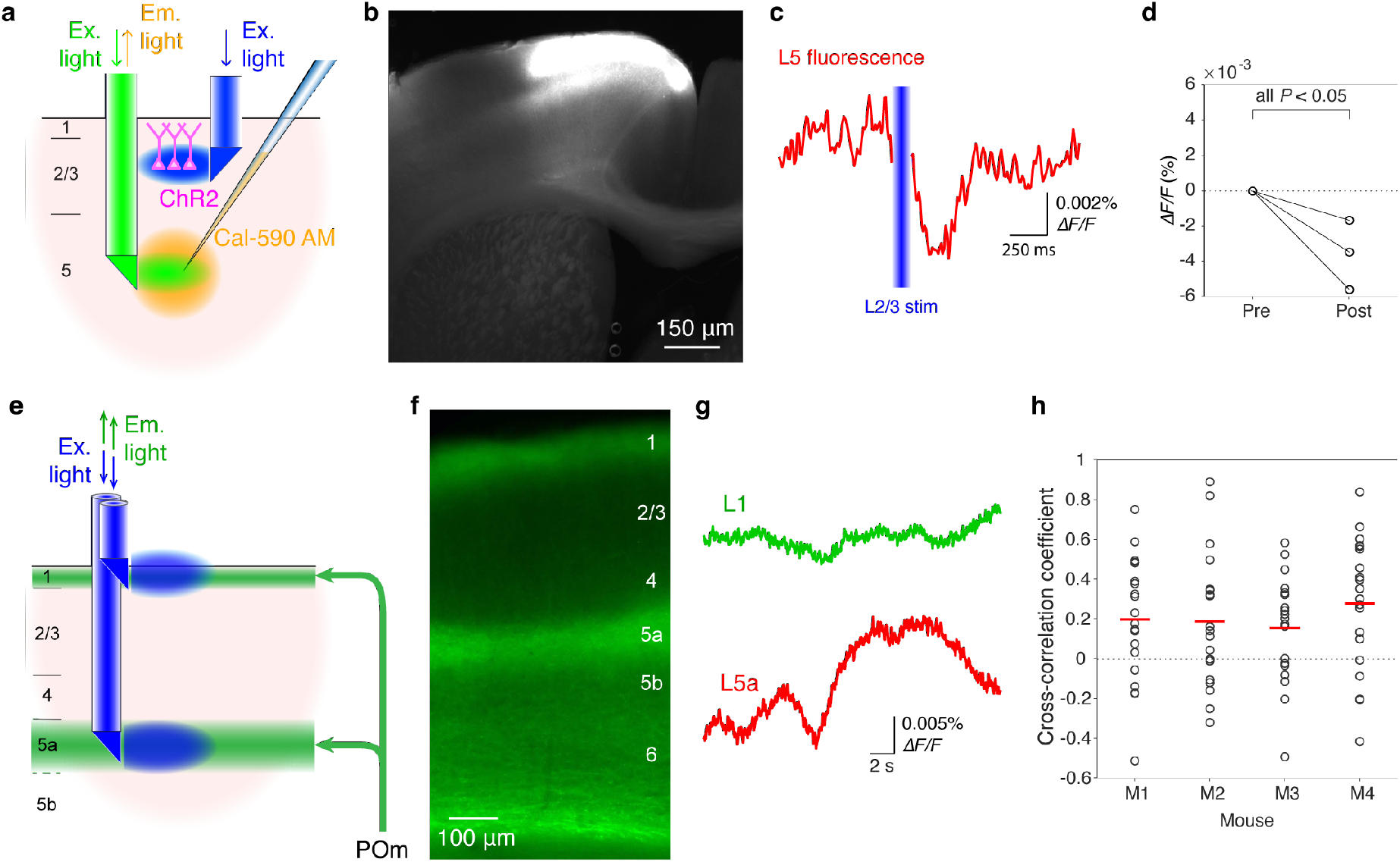
Additional functionalities of double-μPeriscope. (a) Schematic diagram of the experiment that combines optogenetic stimulation and calcium fluorescence imaging. (b) A photomicrograph of cortical slice where L2/3 pyramidal cells express ChR2 and YFP in the primary motor cortex. (c) Fluorescence change in L5 caused by L2/3 stimulation. The trace is the average over 20 measurements. (d) Summary of data from 3 Rasgrf2-2A-dCre mice where the fluorescence reduction in L5 after optogenetic stimulation of L2/3 was statistically significant (n=3 mice, 20 measurements in each mouse, all *P*<0.05, two-tailed student’s t-test). (e) Schematic diagram of the experiment where population fluorescence of thalamic axons distributed in cortical layers 1 and 5a was simultaneously measured. (f) A photomicrograph of cortical slice where axonal terminals from POm are densely distributed in cortical layers 1 and 5a. (g) Example traces of simultaneously measured fluorescence in cortical layers 1 (green) and 5a (red). (h) Summary of cross-correlation coefficients obtained from 4 Gpr26-cre mice. An open circle indicates a pair of simultaneous measurements of axonal activities in layers 1 and 5a. Red line indicates the mean of 20 measurements in each mouse.

Lastly, we demonstrate that a double-μPeriscope can be applied for simultaneous fluorescence imaging of axonal terminals distributed at two depths (Fig 2e). The posteromedial thalamic nucleus (POm) sends axons to cortical layers 1 and 5a in the primary somatosensory area (Fig 2f)^20^; however, it is unknown whether the activities of these two axonal pathways are same or different. Using the double-μPeriscope we measured the axonal activities in cortical layers 1 and 5a simultaneously in the awake head-fixed mice undergoing whisker stimulation. A genetic calcium indicator GCaMP6s^21^ was expressed in POm using a transgenic mouse line Gpr26-cre^22^ and a Cre-dependent adeno-associated virus vector encoding GCaMP6s. We found that the axonal activities in layers 1 and 5a are surprisingly variable; they are clearly uncorrelated in some trials but highly correlated in others (Fig 2g,h). The cross-correlation analysis confirmed this variability, although the mean coefficient was lower than 0.3 in all 4 mice (Fig 2h), suggesting the possibility that POm neurons deliver different types of information to layers 1 and 5a. Further studies are necessary to explain the large variability of axonal activity in layers 1 and 5a. These three sets of preliminary experiments demonstrate that the new tool provides us with a new approach for addressing previously-intractable important neurobiological problems.

Previous studies used an electric lens combined with two-photon microscopy^23–25^ to measure the cellular activities at two depths. The limitations of this approach are 1) that imaging is not simultaneous because the electric shift of focal plane takes time for shifting and stabilization; and 2) that the maximum distance of shift is 500 μm; therefore, the distance between two structures under study must be within this range. A related issue is that imaging deeper structures needs more laser power. Therefore, even if the lens is capable of shifting the focal plane by a larger distance (than 500 μm), the excitation light must be rapidly switched between high and low power in perfect synchronization with the electrical shift of focal plane, which is technically and financially challenging in most laboratories. Besides, deep subcortical structures cannot be imaged with this approach unless a substantial amount of overlaying cortical tissue is surgically removed^26^. The double-μPeriscope we show here does not have these limitations because the distance between two μPeriscopes can be freely adjusted, and imaging is simultaneous no matter how distant the two μPeriscopes are. Deep subcortical structures can also be imaged or optogenetically stimulated with the double-μPeriscope without substantial damage of cortical tissue.

Importantly, the cost to setup the imaging system with a double-μPeriscope is an order of magnitude less expensive. The reason for the significantly lower cost is that it requires only tiny prisms, GRIN lenses and a high sensitivity scientific camera as the main components. Given ever-improving scientific cameras and lowering costs of micro-optics that can now be 3D-printed^27^, approaches using micro-optics such as presented here are likely to be more appealing for many laboratories. We demonstrated its versatility through three sets of experiments where a double-μPeriscope can be used for either optogenetic stimulation, calcium fluorescence imaging, or both of two closely separate cortical layers. The present version of double-μPeriscope uses single-core multimode fiber; therefore, spatial resolution of imaged structure is unavailable. However, a commercially-available multi-core optical fiber bundle would allow this if spatial resolution is required in other applications.

## Methods

### Animal models

Three transgenic mouse lines, Sim1-KJ18-Cre (Tg(Sim1-cre)KJ18Gsat)^28^ × Ai32 (Rosa26-ChR2 reporter mice, JAX 012569)^29^, Rasgrf2-2A-dCre (JAX 022864) and Gpr26-Cre (Tg(Gpr26-cre)KO250Gsat, JAX mouse number 4847098) were used in this study. The age of studied mice was P40-70. Both male and female mice were used. No food or water restriction was imposed. All studied mice were in good health. Mice were housed in single-sex groups in plastic cages with disposable bedding on a 12-hour light/dark cycle with food and water available ad libitum. Experiments were done during both dark and light phase. All procedures were approved and conducted in accordance with the guidelines given by the veterinary office of Landesamt für Gesundheit und Soziales Berlin.

### Virus injection

Rasgraf2-2A-dCre and Gpr26-Cre mice were initially anesthetized with Isoflurane (1%–2.5% in O_2_ vol/vol, Abbott) before ketamine/xylazine anesthesia (75/10 mg per kg of body weight, respectively) was administered intraperitoneally. Lidocaine (1% wt/vol, Braun) was injected around the surgical site. Body temperature was maintained at ~36°C by a heating pad and the depth of anesthesia was monitored throughout the virus injection. Once anesthetized, the head was stabilized in a stereotaxic instrument (SR-5R, Narishige, Tokyo). The skull was exposed by a skin incision and a small hole (~0.5 × 0.5 mm^2^) was made above the primary motor cortex (1.0 mm anterior to bregma and 1.2 mm from midline) of Rasgraf2-2A-dCre mice or the POm (1.8 mm posterior to bregma and 1.25 mm from midline) of Gpr26-Cre mice. AAV9.CaMKII.Flex.hChR2(H134R)-YFP.WPRE3 (Charité Vector Core) or AAV1.Syn.Flex.GCaMP6s.WPRE.SV40 (Addgene #100845) was injected to Rasgrf2-2A-dCre or Gpr26-Cre mice, respectively. Each construct was backloaded into a glass micropipette (Drummond) and was slowly injected (at 20 mL per min, total amount 40-50 nL). The pipette remained there for another 2-5 min after injection. The skin was sutured after retracting the pipette. Rasgrf2-2A-dCre mice were subsequently injected with trimethoprim (TMP, 150 mg per g of body weight) intraperitoneally.

### Fabrication of double-μPeriscope

As described earlier^15^, single μPeriscopes were assembled in-house with a 0.10 × 0.10 mm^2^ or 0.18 × 0.18 mm^2^ micro right-angled prism (Edmund Optics), a 100 μm core multi-mode optical fiber (NA 0.22, Edmund Optics) and a custom-designed Grin lens (NA 0.2, Grin Tech) using a UV curable adhesive (Noland). The two μPeriscopes were precisely aligned with a specific distance between the two prisms and glued with a UV curable adhesive (Noland). Alternatively, two μPeriscopes were not glued but individually manipulated by two stereotaxic micromanipulators (SM-15R, Narishige).

### Extracellular recordings

Animals were initially anesthetized by isoflurane (1-2% in O_2_, vol/vol, Abbott) before urethane anesthesia (0.05 mg per kg of body weight) was administered intraperitoneally. Lidocaine (1%, wt/vol, Braun) was injected around the surgical site. Body temperature was maintained at ~36 °C by a heating pad and the depth of anesthesia was monitored throughout experiment. Once anesthetized, the head was stabilized in the stereotaxic instrument and the skull was exposed by a skin incision. A ~1.0 × 1.0 mm^2^ craniotomy was made above the primary somatosensory (barrel) cortex and the dura matter was removed. The area was kept moist with rat ringer for the entire experiment (135 mM NaCl, 5.4 mM KCl, 1.8 mM CaCl2, 1 mM MgCl2, 5 mM HEPES). A linear array of 16 electrodes (NeuroNexus, A1×16-3mm-50-177-A16) or a glass pipette was perpendicularly inserted into the area such that the uppermost electrode was positioned at 50 μm below the pia. Electrical activity was bandpass-filtered at 1–9 K Hz, digitized at 10 K Hz, amplified by ERP-27 system and Cheetah software (Neuralynx).

### Optogenetic stimulation with a double-μPeriscope

The end of each optical fiber was coupled with a blue LED (peak wavelength 470 nm, Cree or Thorlabs). The timing and intensity of optical stimulation through each μPeriscope was controlled by Power1401 and Spike 2 software (CED) and synchronized with the neural recording system or fluorescence imaging via TTL signals. The light intensity was 12 mW/mm^2^ and the duration was 20 ms. Surgical preparation is same as described in the section “**Extracellular recordings**”. A double-μPeriscope was slowly inserted into the somatosensory cortex such that the upper μPeriscope stimulated the distal apical dendrites (50-150 μm deep from pia) and the lower one stimulated the perisomatic region of L5 pyramidal neurons (500-700 μm deep from pia).

### Calcium imaging with a double-μPeriscope

The imaging setup consisted of a double-μPeriscope, an LED (peak wavelength 470 nm, Thorlabs), an excitation filter (480/30 bandpass, Chroma), an emission filter (535/40 bandpass, Chroma), a dichroic mirror (cutoff wavelength: 505 nm, Chroma), an 80 × 80 pixel high-speed CCD camera with frame rate of 125 Hz (Redshirt Imaging), a 10× infinity corrected objective (58–372, Edmund Optics), and a tube lens (Optem, RL091301-1). Surgical preparation is same as described in the section “**Extracellular recordings**”. An aluminum head implant was fixed to the skull of the mouse with dental cement and the mice were habituated to head-fixation and whisker deflection by a piezo element before imaging. A double-μPeriscope was slowly inserted into the barrel cortex such that the upper μPeriscope covered L1 (0-100 μm deep from pia) and the lower μPeriscope covered L5a (450-550 μm deep from pia). ΔF/F was calculated as (F–F0)/F0, where F is the fluorescence intensity at any time point and F0 is the average intensity over the prestimulus period of 320 ms.

### Calcium imaging combined with optogenetic stimulation

The imaging setup is same as described in the section “**Calcium imaging with a double-μPeriscope**” except an LED (peak wavelength 565 nm, Thorlabs), an excitation filter (555/20 bandpass, AHF), an emission filter (605/55 bandpass, AHF) and a dichroic mirror (cutoff wavelength: 565 nm, AHF). Surgical preparation is same as described in the section “**Extracellular recordings**”. An aluminum head implant was fixed to the skull of the mouse with dental cement and the mice were habituated to head-fixation before imaging. A double-μPeriscope was slowly inserted into the motor cortex so that the upper μPeriscope covered L2/3 (150-300 μm deep from pia) and the lower μPeriscope covered L5 (500-700 μm deep from pia). The calcium indicator Cal-590 AM (AAT Bioquest) was backloaded into a micropipette (Drummond) and slowly injected (at 20 nl per min, total 40–50 nl) to L5 (~600 μm below pia) of the primary motor cortex 1.5–2 h before imaging experiments. The pipette remained there for at least 5 min after injection. ΔF/F was calculated in the same way as described in the section “**Calcium imaging with a double-μPeriscope**”.

### Data analysis and statistical methods

Analyses were conducted using Matlab (Mathworks). Significance was determined by two-tailed, paired student’s t test at a significance level of 0.05. The statistical test was chosen based on the data distribution using histogram. No statistical method was used to predetermine sample sizes, but our sample sizes are those generally employed in the field. We did not exclude any animal for data analysis. The variance was generally similar between groups under comparison. No blinding/randomization was performed.

## Notes

### Competing Interest Statement

The authors have declared no competing interest.

